# Recording action potential propagation in single axons using multi-electrode arrays

**DOI:** 10.1101/126425

**Authors:** Kenneth R. Tovar, Daniel C. Bridges, Bian Wu, Connor Randall, Morgane Audouard, Jiwon Jang, Paul K. Hansma, Kenneth S. Kosik

## Abstract

The small caliber of central nervous system (CNS) axons makes routine study of axonal physiology relatively difficult. However, while recording extracellular action potentials from neurons cultured on planer multi-electrode arrays (MEAs) we found activity among groups of electrodes consistent with action potential propagation in single neurons. Action potential propagation was evident as widespread, repetitive cooccurrence of extracellular action potentials (eAPs) among groups of electrodes. These eAPs occurred with invariant sequences and inter-electrode latencies that were consistent with reported measures of action potential propagation in unmyelinated axons. Within co-active electrode groups, the inter-electrode eAP latencies were temperature sensitive, as expected for action potential propagation. Our data are consistent with these signals primarily reflecting axonal action potential propagation, from axons with a high density of voltage-gated sodium channels. Repeated codetection of eAPs by multiple electrodes confirmed these eAPs are from individual neurons and averaging these eAPs revealed sub-threshold events at other electrodes. The sequence of electrodes at which eAPs co-occur uniquely identifies these neurons, allowing us to monitor spiking of single identified neurons within neuronal ensembles. We recorded dynamic changes in single axon physiology such as simultaneous increases and decreases in excitability in different portions of single axonal arbors over several hours. Over several weeks, we measured changes in inter-electrode propagation latencies and ongoing changes in excitability in different regions of single axonal arbors. We recorded action potential propagation signals in human induced pluripotent stem cell-derived neurons which could thus be used to study axonal physiology in human disease models.

**Significance Statement:** Studying the physiology of central nervous system axons is limited by the technical challenges of recording from axons with pairs of patch or extracellular electrodes at two places along single axons. We studied action potential propagation in single axonal arbors with extracellular recording with multi-electrode arrays. These recordings were non-invasive and were done from several sites of small caliber axons and branches. Unlike conventional extracellular recording, we unambiguously identified and labelled the neuronal source of propagating action potentials. We manipulated and quantified action potential propagation and found a surprisingly high density of axonal voltage-gated sodium channels. Our experiments also demonstrate that the excitability of different portions of axonal arbors can be independently regulated on time scales from hours to weeks.

## Introduction

Central nervous system (CNS) axons are the output structures of neurons, integrating subthreshold synaptic potentials and translating that to an all-or-none digital event through axonal action potential propagation. In spite of this central role in information transfer, the physiology of CNS axons has not been studied is as much detail as dendrites or neuronal cell bodies. Though their small caliber likely leads to their relative inaccessibility, several groups have used either paired axonal recordings to study the fidelity of propagation (Meeks et al., 2005; Khaliq and Raman, 2005; Raastad and Shepherd, 2003), and the density of voltage-gated sodium channel (Hu and Jonas, 2014). Others have used voltage-sensitive dyes to investigate the heterogeneity of electrophysiological characteristics in different types cortical interneurons (Casale et al., 2015). Technically challenging experiments such as these demonstrate the difficulty of experimentally accessing CNS axons.

MEAs are tools for recording eAPs (also referred to as “spikes”) from hundreds of neurons simultaneously (Lewis et al., 2015; Liu et al., 2012). When neurons and glia are cultured on planer MEAs, the electrical behavior of self-organized neural ensembles can be non-invasively monitored over days to weeks (Potter and DeMarse, 2001). In these cultures, close apposition of neurons with each recording electrode can produce recordings with high signal-to-noise characteristics, while the ease of experimental manipulations in cell culture systems creates opportunities for experiments that might otherwise be technically challenging *in vivo*. Because outgrowth of neuronal processes and electrode orientation are in the same plane, it is not unreasonable to expect that action potential propagation within single neurons might be detected across multiple electrodes in low electrode density arrays.

We report here eAPs at groups of electrodes with characteristics expected of the detection of action potential propagation at multiple sites. This includes the repeated and consistent co-occurrence of spikes from cultured mouse hippocampal neurons on multiple MEA electrodes. Within co-active electrode groups, eAPs occurred with invariant sequence and with inter-electrode latencies that are consistent with propagation velocities reported for unmyelinated axons (Kress and Mennerick, 2009). Co-active electrode groups were present in all our data records and recordings from each unique culture often displayed multiple distinct and independent co-active electrode groups. Repeated eAP detection by multiple electrodes confirms each coactive set of spikes originates from single neurons and the stereotyped sequence and pattern at co-active electrodes unambiguously fingerprints each neuronal source of these signals. These unambiguously identified neurons can be used to gauge the behavior of single neurons within the ensemble. Averaging the eAP co-occurrences across all MEA electrodes of each group of co-active electrodes revealed electrodes with subthreshold eAPs, increasing the two-dimensional extent of propagation and thus the temporal range of inter-electrode latencies. We found that the fidelity of action potential propagation is unaffected even when we significantly decrease the density of voltagegated sodium channels, consistent with high axonal sodium channel density. We simultaneously measured both increases and decreases in eAP amplitudes in different regions of the same axonal arbor occurring over several hours. Over longer time periods, the extent of propagation of single axonal arbors dynamically expands and contracts over multiple weeks. Recording over longer time periods also revealed changes in spike waveforms consistent with decreasing transmembrane action potential duration, changes in the extent and duration of repolarizing conductances and time-dependent changes in the speed of propagation. We also detected these signals in cultures of human iPS-derived neurons, creating the potential for the study of axonal pathophysiology in human genetic conditions.

We used these propagation signal eAPs to reveal the details of axonal excitability from single axons during neuronal network development in an experimentally amenable *in vitro* system. Multi-site, non-invasive extracellular recording can be used to study axonal physiology in the axonal arbors of unambiguously identified single neurons within the larger neuronal ensemble and to monitor how the properties of action potentials in various portions of axonal arbors change over time.

## Methods

### Cell culture

Cleaned multi-electrode arrays (MultiChannel Systems; 120MEA100/30iR-ITO arrays) were sterilized with UV irradiation (for ∼30 minutes), then incubated with a poly-D- or poly-L-lysine (0.1 mg/ml) solution for at least one hour, rinsed several times with sterile de-ionized water and either plated immediately after rinsing or allowed to dry before cell plating. The culture chamber surrounding the MEA (19 mm inner diameter) was filled with 1 mL of cell culture media. To allow enough time for glia to proliferate and become confluent in the area around the electrodes, cell cultures were prepared in two stages. The first plating was intended to seed the MEA substrate with proliferating glial cells and the second plating was intended for neurons to grow upon a confluent glia substrate. Cells were plated at 100,000 to 125,000 cells per dish for the first plating and at 125,000 to 200,000 cells per dish for the second plating. Mouse hippocampal neurons were used for most of the experiments described here. All mice were from a C57BL/6 genetic background and male mouse pups were used for all cell cultures. For cell culture, mouse pups were decapitated at P0 or P1, the brains were removed from the skulls and hippocampi were dissected from the brain (Tovar and Westbrook, 2012). After the first plating most neurons did not survive. However, when necessary for timing purposes, cultures were treated with 200 μM glutamate for 30 minutes at 37° C to kill any remaining neurons. This was done 5-7 days after the first plating. Cultures were grown in a tissue culture incubator (37°C, 5% CO2), in a medium made with Minimum Essential Media with 2 mM Glutamax (Life Technologies), 5% heat-inactivated fetal calf serum (Life Technologies), 1 ml/L of Mito+ Serum Extender (BD Bioscience) and supplemented with glucose to an added concentration of 21 mM. Cell culture media was changed 2 times per week. Cultures were maintained for as long as 6 weeks. No antibiotics were used in our cell culture system. All animals were treated in accord with University of California and NIH policies on animal care and use.

### Solutions, electrophysiology and analysis

The recordings described in this work were done in cell culture medium (see above) so as to minimally disturb the neurons and maintain sterility. In some cases we instead used an extracellular solution containing (in mM) 168 NaCl; 2.4 KCl; 10 HEPES; 10 D-glucose; 1.8 CaCl2; and 0.8 mM MgCl2. The pH was adjusted to 7.4 with KOH. The osmolality of external and internal solutions was adjusted to 320 mosmol. Salts were obtained from Sigma-Aldrich or Fluka; some drugs (TTX, NBQX, CPP, gabazine) were obtained from Ascent Scientific. For the experiments reported here, drugs were introduced directly into the recording chamber.

Recordings were done using MultiChannel Systems MEA 2100 acquisition system. Data were sampled at 20 kHz and post-acquisition bandpass filtered between 200 and 4000 Hz. Data were recorded on all 120 data channels. We controlled the head stage temperature with an external temperature controller (MultiChannel Systems TC01). Most recordings reported here were done at 30^°^ C, unless otherwise indicated. In our conditions, the working ambient head stage temperature was ∼29 ^°^ C and a small subset of early experiments were done without temperature control. The ambient head stage temperature set the minimum temperature at which we could perform temperature experiemnts.

All recordings were done on neurons at 5-30 days *in vitro* (DIV). We only used recordings with signals present on the majority of channels. Recordings were typically 3 to 5 minutes long. Recording duration was kept short to minimize the effects of removing MEAs from the incubator and to minimize data file size. To minimize the effects of evaporation, maintain cell culture sterility and decrease degassing of the media, all recordings were done with the MEAs covered by a CO2-permeable, water vapor-impermeable membrane (Potter and DeMarse, 2001) which was held in place over the recording chamber by a Teflon collar placed over the culture chamber. Following placement of each MEA in the recording head stage, each array was allowed to equilibrate to head stage temperature (requiring at least 5 minutes while we monitored the re-equilibration at the recording stage to the set temperature) prior to recording. Recording durations were minimized to avoid large changes in CO2 and pH. Though our recordings were done with covers over the culture reservoir surrounding the MEA, separate control experiments showed that minimizing the recording duration was important because the temperature gradient between the temperature control element and the top of the MEA chamber lid promoted evaporation from the small volume of the culture wells and sharply increased the osmolarity of the culture media (data not shown). For experiments requiring temperature changes, head stage temperature was monitored and each MEA was kept at the new temperature for at least 5 minutes.

### Spike Detection and Analysis

MultiChannel Systems proprietary files were converted to HDF5 file format prior to all analysis. Extracellular signals were bandpass filtered using a digital 2nd order Butterworth filter with cutoff frequencies of 0.2 and 4 kHz. Spikes were then detected using a threshold of 6 times the standard deviation of the median noise level (Quiroga et al., 2004). No spike sorting was done in any of our experiments. Analysis was done using Igor (Wavemetrics) and with custom software written in Python and Mathematica. In brief, for each eAP detection time, 5 data points centered on the minimum spike time are fit to a 2nd order polynomial whose minima is calculated to provide an estimate of the precise spike time. A 3 ms window of the extracellular recording was then extracted for each spike time. Action potential propagation signals were initially detected by eye with the help of custom spike visualization software and validated by signal averaging using custom software. We subsequently developed software to automate propagation signal detection based on repeated co-occurrence of spikes on stereotyped groups of electrodes. The criteria for automated detection was that these spikes must occur with high incidence and within a specified inter-electrode time window. All statistical data are displayed as the mean ± standard deviation. For the cases of multiple comparisons, we used analysis of variance followed by the Bonferroni multiple comparison correction. Linear fits of data from manipulating propagation latency were done using Igor. The percent change in propagation velocity was calculated by:

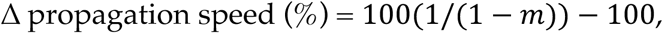
where *m* is the average slope for each condition (temperature or TTX concentration).

### Immunocytochemistry

For immunocytochemistry experiments, the cells were grown on coverslip glass in plastic culture dishes but otherwise prepared in the same manner as neurons plated on MEAs. Cells were fixed with 4% paraformaldehyde for 15 min at room temperature and washed 3 times with phosphate-buffered saline (PBS). Cells were then permeabilized with 0.25% Triton X-100 in PBS for 10 min at RT and washed 3 times with PBS. Cells were blocked with BlockingAid (Life Technologies) for 1 hour at room temperature and incubated with primary antibodies in the block solution at 4° C overnight. Primary antibodies were diluted at 1:2000 for the guinea pig anti-vGluT1 (Synaptic Systems #135304) and at 1:1000 for the rabbit anti-GAD67 (Synaptic systems #198013). The cells were washed 3 times with PBS and incubated with the secondary antibodies in the blocking solutions for 1 hour at room temperature. The donkey anti-guinea pig conjugated with Alexa-647 (Emd Millipore #AP193SA6) and donkey anti-rabbit conjugated with Alexa-488 (Life Technologies #A21206) secondary antibodies were diluted at 1:1000. Cells were washed with PBS 3 times and mounted on glass slides with ProLong Gold Antifade Mountant (Fisher #P36934).

### Human iPS-derived Neurons

Neurons were derived from human induced pluripotent stem cells (iPSCs) by NeuroD1 overexpression as described previously (Lalli et al., 2016) Briefly, hiPSCs were passed as single cells with Accutase (StemCell Technologies) and were plated on Matrigel-coated (Corning) plates in mTeSR1 (StemCell Technologies) with ROCK inhibitor (Y-27632, StemCell Technologies). The following day, cells were transduced with lentiviral vector NeuroD1-GFP-Puro driven by a tetracycline inducible promoter. Two days after transduction, doxycyclin (1μg/ml) was added to the media to induce the transgene, and was designated as Day 0. Transduced cells were selected on Day 1 by adding puromycin (1μg/ml). On Day 3, cells were lifted with Accutase and replated on MEAs with mouse glial cells already seeded. From Day 5 to Day 8 half media change was performed with doxycyclin and AraC (1μM) to kill the proliferating cells in the culture. The induced neurons were maintained and recorded in N2/B27 media.

## Results

### Identifying action potential propagation

In recordings from each culture of mouse hippocampal neurons on MEAs, we consistently noticed groups of electrodes that were repetitively co-active. Figure 1A shows the electrode layout of the MEAs used in our experiments, and the position of one such group of co-active electrodes indicated (black circles). Electrode pitch was 100 microns, center-to-center and each electrode was 30 microns in diameter. An example of repeated spike co-occurrence from six electrodes is shown in Figure 1B. The earliest occurring spike in this sequence was always at electrode F9 and the last spike was at electrode D4. An example of repeated eAP co-occurrence from these six electrodes is shown (arrows, Figure 1B). For the 180-second-long data record, spikes repeatedly cooccurred (14.6 Hz) on these electrodes. The spike sequence during each co-occurrence event was invariant among co-active electrodes. The invariant sequence stereotypy and the latency in spike timing between each constituent electrode is shown (Figures 1C). Each panel shows the average waveform of 200 spikes from each indicated electrode (black traces) superimposed on 10 individual traces from that electrode (thin grey traces). In this example, eAP waveforms from other electrodes were indexed to the negative peak of eAPs at electrode F9. Spikes at electrode F9 had the highest amplitude in this group of co-active electrodes. The absence of an initial upward capacitive component of the spike waveform at electrode F9, the amplitude relative to eAPs at other co-occurring electrodes and the fact that eAPs at electrode F9 occur before eAPs at other co-occurring electrodes is consistent with these eAPs originating at or near the axon initial segment (AIS). The characteristic delay between spikes and the sequence stereotypy of these signals during each co-active event is apparent from the averaged events in this figure.

**Figure 1.**
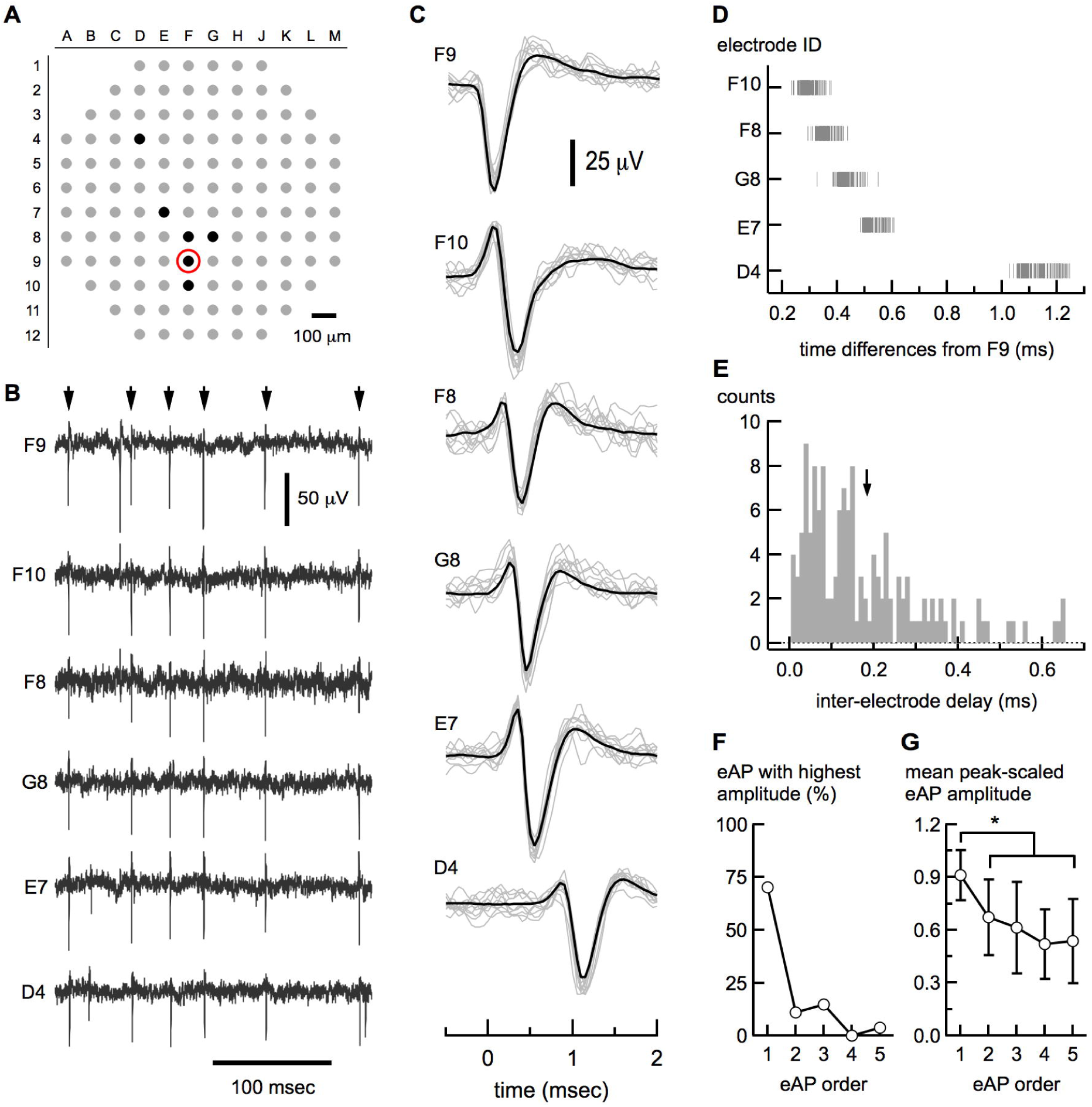
Co-active electrodes reveal action potential propagation. A map of the extracellular electrode configuration (A) is shown. Electrode pitch spacing is 100 microns, center-to-center and electrodes are 30 microns in diameter. The black circles indicate repeatedly co-active electrodes. The earliest spike in the repeating sequence always occurred at F9 and terminated at D4. The red circle indicates electrode F9. The extracellular voltage records (B) from the 6 electrodes indicated (A) are shown; arrows highlight co-occurring spikes amongst these electrodes. Electrode designations are to the left of each trace. Spikes from co-active electrodes are shown at higher time resolution (C). Averages of 200 spikes (thick black lines) are shown superimposed on 10 individual sweeps (grey lines). Electrode designations are indicated in the upper left. Note the increasing delay with distance from F9. The difference in spike time peaks between spikes at F9 and all other electrodes of this co-active sequence are shown (D). The high coherence, very short time delays and low variability (CVs from 0.047 to 0.080) are consistent with action potential propagation. The inter-electrode timing differences between 139 electrode pairs is displayed in (E). The mean (± SD) difference (arrow, 0.18 ± 0.15 ms) is much shorter than expected for direct synaptic coupling between neurons. In 28 propagation signals with 5 component electrodes, the electrode with the on average the electrode with the highest amplitude was the first electrode in the series (F). The mean amplitudes (± SD) for spikes in 5-component propagation signals are plotted (F). Spike amplitudes were normalized to spikes within each propagation signal. In this group of propagation signals, 19/28 of the highest amplitudes were recorded from the first electrode, consistent with the high voltage-gated sodium channel density at the site of action potential initiation.

Detection of eAPs at the same neuronal position by two electrodes can occur in cases where electrodes are closely apposed or, theoretically, in cases where signal amplitude is quite large. However, such signals are expected to have similar waveform and to be detected simultaneously, unlike the distinct eAP waveforms from F9 and F10 (Figure 1C) which occurred with a consistent delay between electrodes. This is contrary to the expectation of simultaneous detection by neighboring electrodes but is expected for cell-intrinsic signal propagation. This slight spike but consistent latency between electrodes was the norm in the vast majority of the cases we examined in detail. The morphological variability in spike waveforms between electrodes of co-active groups is also inconsistent with signal bleed-through between electrodes. The combined effect of the large electrode pitch (100 microns) and the fact that extracellular voltage decreases steeply (as 1/r^2^; Buzsaki et al., 2012) likely limits simultaneous eAP detection by multiple electrodes in our assays.

If eAPs at co-active electrode groups represent action potential propagation, we would expect the spike latencies between electrodes to be consistent with reported measurements of propagation from unmyelinated axons. Figure 1D shows the latencies between spikes at different component electrodes within the first 250 co-active spikes in the electrodes from Figure 1B. Hash marks indicate the delay between spikes at the index electrode (F9) and the indicated electrodes for each incidence of co-activity. In the example from Figure 1C, the signal at electrode F9 propagates to electrode D4 (539 microns distant) at ∼0.5 m/sec. On average, the propagation rate between electrodes separated by at least 300 microns was 0.68 ± 0.32 m/sec (n = 42). These values assume a direct path between electrodes and thus likely underestimate the action potential propagation velocity. However, these estimates are consistent with reported measurements of action potential propagation in unmyelinated axons from hippocampal neurons (Meeks and Mennerick, 2007; Hu and Jonas, 2014). Additionally, the coefficient of variation (CV) of the latencies between spikes at F9 and spikes at each co-active electrode were low, ranging from 0.047 to 0.080, consistent with a high fidelity process like action potential propagation (Meeks et al., 2005). We plotted the distribution of time differences between consecutive spikes from 62 groups of co-active electrodes in 6 recordings from unique cultures (Figure 1E). The mean inter-electrode interval between consecutive spikes from this group was 0.18 ± 0.15 ms (n = 139), much shorter than expected for direct synaptic coupling between neurons (3 - 5 ms; Ivenshitz and Segal, 2010). In our example, the mean sequential inter-electrode latencies, which ranged from 0.064 ms to 0.59 ms, were much shorter than the median inter-spike interval (19.7 ms) from all the electrodes of this MEA. Combining this with the number of co-occurrences (2635) and the coefficient of variation of the latencies makes it unlikely that these repeated sequences resulted from chance. Latencies of eAPs among co-active electrode groups on this scale and the low spike timing variability between coactive electrodes is consistent with these groups of electrodes detecting action potential propagation in different parts of the same neuron.

Action potential initiation occurs at the AIS (Bean, 2007; Kole and Stuart, 2012), the site on neurons with the highest density of voltage-gated sodium channels (Kole et al., 2008; Hu et al., 2009). If the co-active electrode groups in our recordings reflect action potential propagation, then, other things being equal, we might expect that the initial spike in co-active electrode sequences would tend to be the largest. We tested whether we could detect a correlation between eAP amplitude and the order of occurrence within 28 unique groups of co-active electrodes from several cultures. Each group was composed of 5 co-active electrodes. For these groups, we first determined the eAP sequence, then peak-scaled the eAP amplitudes of each group to the amplitude of largest average spikes within that group. The eAP with highest amplitude usually occurred (70.4% of the time) at the first electrode in each sequence (Figure 1F). We averaged the eAP amplitude at each position to get the mean peak-scaled eAP amplitude for each of the five electrodes of the sequence. As seen (Figure 1G), the first spike in these sequences, on average, had the highest peak-scaled amplitude (0.91 ± 0.14; p < 0.00001; ANOVA and *post hoc* Bonferroni correction). The robustness of these results is surprising because the amplitude of the extracellular voltage signal decreases steeply with distance from the source (Anastassiou et al., 2015) and the distance between cellular component and electrode is a random unknown variable. That the initial spike in co-active electrode sequences, on average, had the largest amplitude is consistent with detection of signals at or near the AIS. The lack of an initial capacitive (upward) peak in the first eAP of many of these sequences is also consistent with eAP detection at or near the AIS because eAPs detected at the site of action potential initiation are not expected to be contaminated by stray capacitance due to signal propagation. These data also indicate that the transmembrane conductance is decreased by only ∼42% at sites distant from the initiation site, supporting the idea that these signals are from axons. The repeated eAP co-occurrence and inter-electrode latencies support the idea that spikes at these electrode groups represent action potential propagation at different cellular regions. The density of voltage-gated sodium channels at the AIS and in axons is many times higher than in the soma or dendrites (Hu et al., 2009; Lorincz and Nusser, 2010; Cox et al., 2000). This steep gradient of voltage-gated sodium channels in neurons makes it likely that signals from co-active electrode groups represent detection of axonal action potential propagation, in contrast to action potential back-propagation into soma or dendrites. Because the eAPs among co-active electrodes have properties consistent with action potential propagation, we refer to the collection of eAPs as propagation signals.

### Propagation signals are widespread

In recordings from mouse neurons, propagation signals were seen in all our data records, with a mean of 7.5 ± 3.4 signal per array from 75 randomly chosen arrays (Figure 2A). From this same data set, the distribution of constituent electrodes per propagation signal had an average of 4.1 ± 2.4 electrodes per signal (Figure 2B). Thus on average, spike redundancy due to propagation signals affected roughly 25% of the MEA electrodes in every recording. To demonstrate the impact of propagation signal redundancy on the raw data record, we plotted the eAP time stamps of 6 electrodes from a single MEA recording, showing spike time before (black hashes) and after (red hashes) removing propagation signals. In these records, propagation signals constitute anywhere from 18.9% (electrode H1) to 87.0% (electrode J1) of the total number of spikes from these electrodes (Figure 2C). In 25 unique recordings, removing propagation signal eAPs from all but one component electrode in each array decreased the total number of detected spikes on these MEAs by 41.2 ± 19.2%. We tested how the electrode pitch in our experiments (100 microns) contributed to the frequency of propagation signal detection by retrospectively removing the contributions from every other electrode in data from 100 micron pitch arrays. We used propagation signals with 4 super-threshold electrodes (the mean number of electrodes per propagation signal) and still detected propagation signals in 12 of 39 cases, indicating that larger interelectrode distances did not eliminate this form of signal redundancy. Propagation signals are present in every recording from cultured mouse neurons and most recordings contained multiple unique examples meaning that they could provide a model to study axonal physiology at multiple sites of single neurons. However, the significant signal redundancy produced by propagation signals could compromise spike rate analysis.

**Figure 2.**
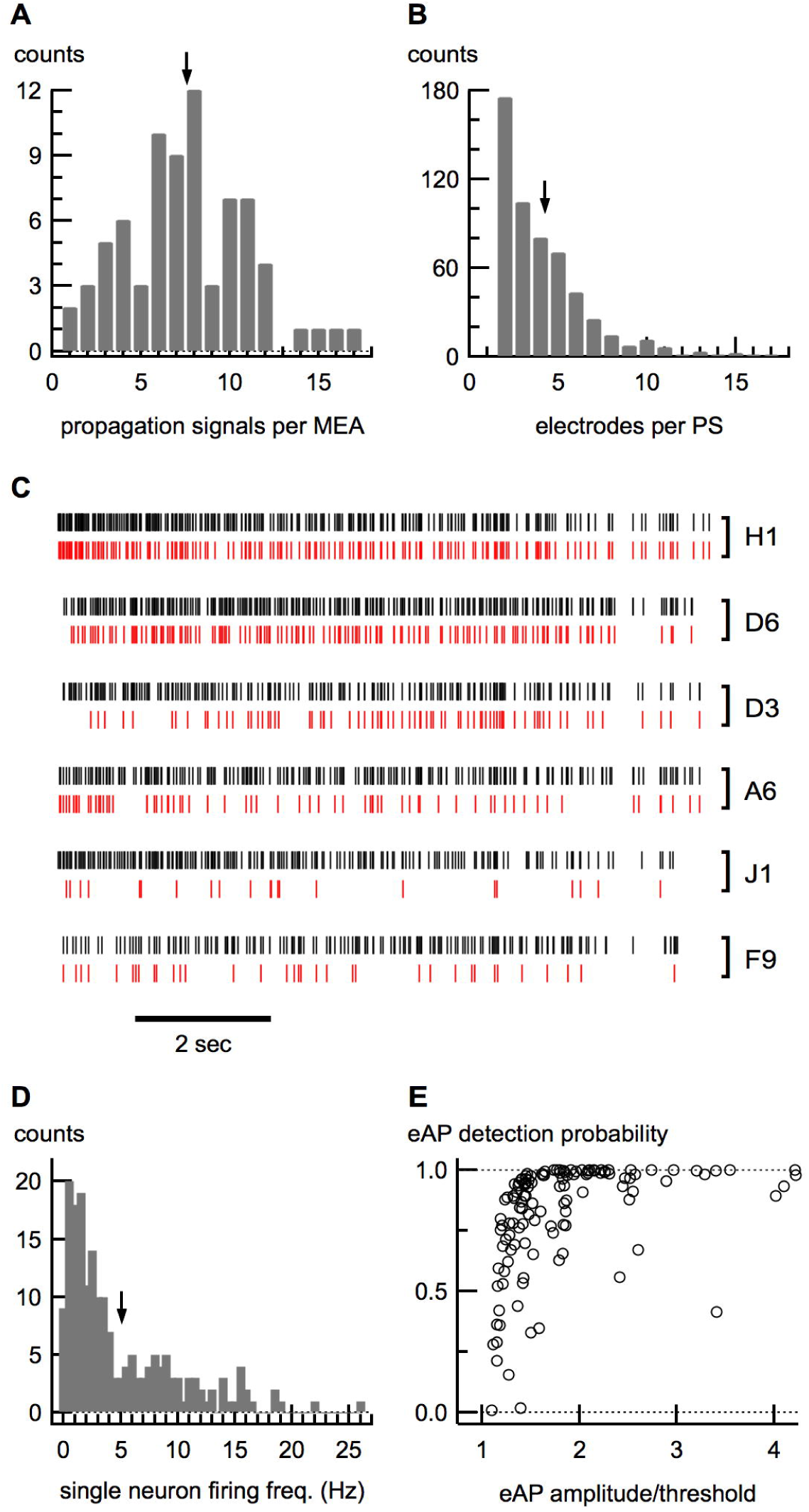
Widespread occurrence of propagation signals. In a distribution of the number of propagation signals per MEA from 75 unique recordings (A), the average number of propagation signals per MEA (± SD) was 7.5 ± 3.4. The distribution of electrode components per propagation signal is shown in B. For 544 propagation signals, the average number of electrodes (± SD) was 4.1 ± 2.4. The effect of removing propagation signals on spike train analysis is shown (C). Ten seconds of spike train data from 6 electrodes from the same MEA recording are shown. Spike trains before (black) and after (red) removal of propagation signal removal are shown. Electrode designations are shown to the right of each spike train pair. These data were not spike sorted and demonstrate that removal of propagation signals is itself a spike sorting step. The distribution of neuronal firing frequency from 192 propagation signals is shown (C) with the mean (± SD) of 5.03 ± 5.02 Hz indicated by the arrow. For several propagation signals, the ratio of detecting spikes at other constituent electrodes compared to spikes in the initial electrode is expressed as a probability and is plotted against the ratio of the eAP amplitude to the detection threshold for that electrode (D). This plot shows that as the effective signal-to-noise ratio of individual spike components increases so does the detection probability. Arrows in A, B and D indicate the means of each indicated distribution.

The nature of extracellular recording precludes knowing the neuronal source of any set of eAPs. However, because propagation signals represent action potentials measured in different regions of single neurons, the repeated co-detection by multiple electrodes identifies that these spikes result from single neurons. The eAP sequence and spatial arrangement of electrodes in each propagation signal creates a ‘fingerprint’ that identifies each signal. Thus we can monitor the firing frequency of single identified neurons in the background of eAPs from neurons at all other electrodes. The neuronal firing frequency distribution of 192 propagation signals from 39 unique MEA records is shown (Figure 2D). This distribution of firing frequencies is consistent with other reports of action potential frequency from hippocampal neurons *in vivo* (Ranck, 1973; Fenton and Müller, 1998). The frequencies at which these neurons fire action potentials provide enough co-occurring eAPs for signal averaging and thus a significant increase in the signal-to-noise ratio of propagation signal eAPs.

In many propagation signal sequences there were instances when spikes at constituent propagation signal electrodes were occasionally not reliably detected, even when other spikes of the same co-active electrode group were detected. This could represent failures of action potential propagation, as might be expected in cases of failures to invade axonal branches. Alternatively, this could result from signal detection failures of signal detection in cases when spike amplitude is near the detection threshold. We examined the source of failures by plotting the probability of eAP detection at constituent propagation signal electrodes as a function of the ratio of eAP amplitude to our spike detection threshold. As shown (Figure 2E) the probability of spike detection tended to decrease with the spike height/threshold ratio, indicating that the occasional inability to detect component propagation signal eAPs at constituent electrodes likely result from signal detection issues rather than compromises in action potential propagation. This result is not surprising given the reported high reliability of action potential transmission in hippocampal axons (Cox et al., 2000; Raastad and Shepherd, 2003) but does speak to the level of signal interference from background recording noise. Our data show that propagation signals are widespread in all our cultures and occur with high enough frequency for signal averaging. However, the high incidence of propagation signals could significantly interfere with spike train analysis.

### Action Potential Propagation in Single Neurons at Multiple Sites

Repeated detection of spikes at multiple co-active electrodes validates that these signals originate from single neurons and can thus be averaged. As shown in Figure 3A, signal averaging reveals a greater number of constituent propagation signal electrodes because of the decrease in the signal-to-noise ratio in the averaged record from each electrode. In this example, consistent co-activity of electrodes D9 and F9 was used as the basis for isolating this propagation signal. We recorded more than 1900 co-occurring spikes in 180 seconds of recording. We averaged 800 of these co-occurring spikes at all MEA electrodes from this recording using a 5 millisecond window centered around the first spike (D9) of the co-occurring pair. Signal averaging uncovered electrodes with eAPs that were otherwise below the detection threshold. The additional electrodes increased the cohort of inter-electrode latencies for each neuron and revealed a much larger two-dimensional extent of action potential propagation. For the example propagation signal (Figure 3A), signal averaging revealed eAPs at 14 additional electrodes. Averaged eAP waveforms (red traces) are superimposed on 10 sweeps of raw data (black traces) in a subset of these electrodes (Figure 3A). The data from each electrode show that most signal averaged eAPs were well below the detection threshold (indicated by the grey band in each data window). The similarity in waveform morphology of super- and sub-threshold eAPs is shown by superimposing the peak-scaled averaged waveforms of 2 super-threshold spikes and 2 sub-threshold spikes (Figure 3B). In this example with just 2 super-threshold spikes, the inclusion of subthreshold eAPs demonstrates the latency between eAP peaks and doubles the range of inter-electrode latencies between the index electrode (D9 in this case) and all other propagation signal electrodes. In a subset of 47 unique propagation signals, the average number of electrodes with super-threshold spikes was 3.4 ± 1.6 per propagation signal. Signal averaging each of these propagation signals increased the number of electrodes with eAPs to 18.4 ± 9.2, or roughly 15% of the total number of MEA electrodes. By increasing the apparent two-dimensional extent of action potential propagation, signal averaging eAPs based on spike co-occurrence also increases the temporal range of propagation latencies among the sampling electrodes.

**Figure 3.**
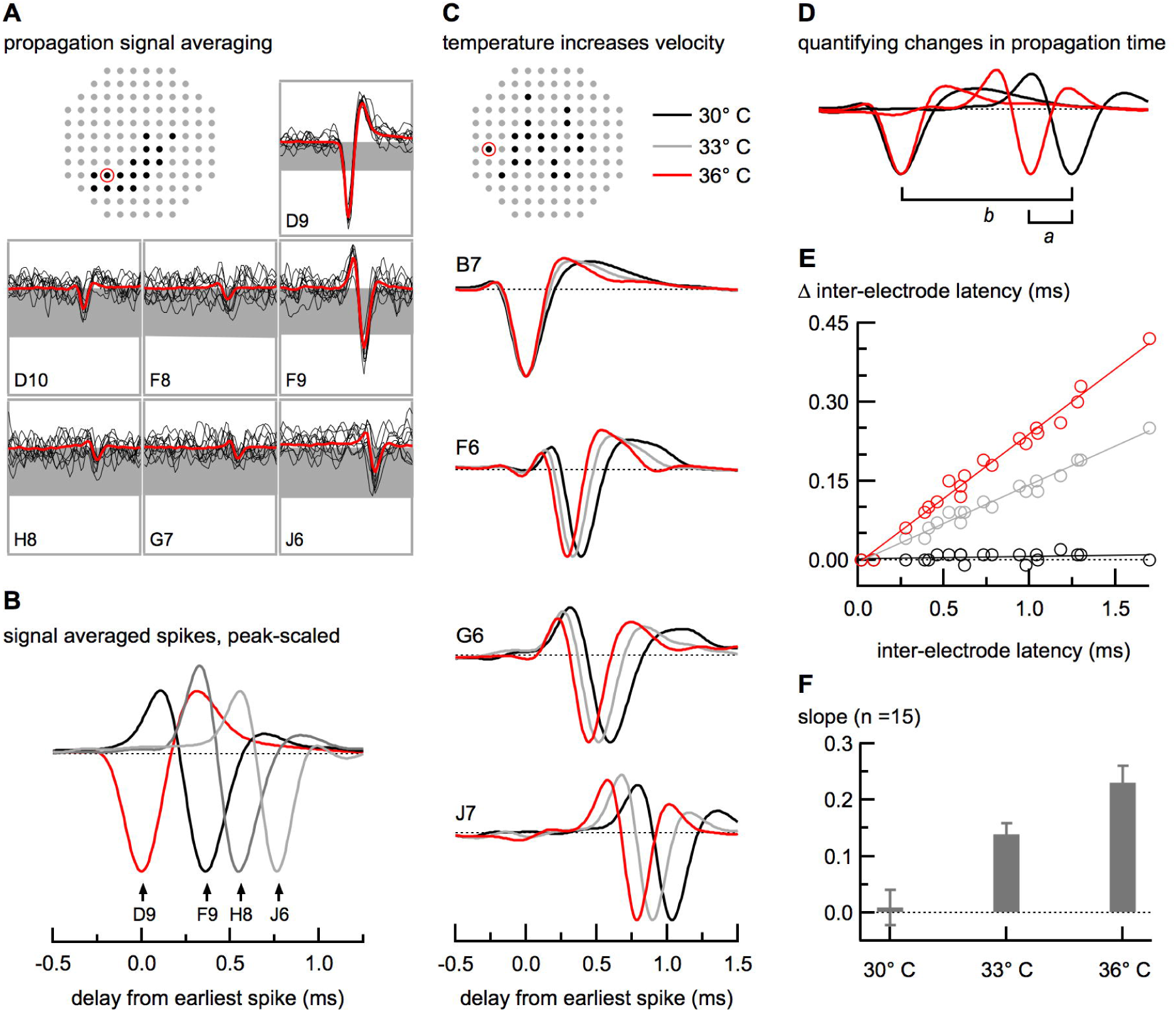
Using signal averaging to study action potential propagation. A map of propagation signal component electrodes, with sweeps from a subset of those electrodes, centered around the spike at D9 is shown (A). The spike detection threshold for each electrode is indicated by the grey band. Data from each electrode shows a signal averaged eAP (red) super-imposed on ten raw data traces (black). For this signal, electrodes D9 and F9 consistently had spikes above the detection thresholds; eAPs at other electrodes were only revealed by signal averaging. Vertical axes were the same for all electrodes (50 μV) except for for D9 (90 μV). Horizontal axes were the same for all electrodes (3 ms). A subset of these super- and sub-threshold eAPs was peak-scaled to the largest negative peak (from electrode D9) and super-imposed to show propagation delays between constituent electrodes (B). Spikes at more distant electrodes occurred later in the sequence. Sub-threshold eAPs are those that are revealed after signal averaging. A map of electrodes showing super- and sub-threshold eAPs for another propagation signal, along with a subset of those eAPs is shown (C). Recordings were done at 30, 33 and 36° C. As expected, increased temperature decreased the latency between electrodes. The shorter time intervals between the initial spike (at electrode B7) and spikes at other electrodes is evident in the leftward shift in time of these eAPs. Temperatures are indicated by color and eAP waveforms were peak-scaled. Because action potential propagation is expected to be linear, experimentally induced changes in propagation velocity should also be linear. For each propagation signal, we assessed changes in propagation velocity by plotting the change in the latency (‘*a*’ in Figure 3D) between the index (first) eAP and all other eAPs of this propagation signal as a function of the latency between the initial spike and each spike time in control conditions (‘*b*’ in Figure 3D). The temperature-dependent increase in propagation velocity is seen as an increase in the slope of the linear fit of the data. Super- and sub-threshold signals were combined for this analysis. Data from control experiments is shown on the horizontal 0 line. For this example, the data was well-fitted with a straight line (r^2^=0.986). Spikes in Figure 3C, D and E were peak-scaled to the largest negative peak. Aggregate temperature data for each condition is shown (F). Data are presented as mean ± SD. Red circles in A and C indicate the index electrode for the indicated propagation signal.

Measurement of action potential conduction velocity requires knowing the propagation path length. Obtaining this information is routinely impractical in our system due to the high density of cells (and thus processes) in our neuronal cultures. To quantify the effects of experimental manipulations known to affect action potential propagation, we plotted the change in inter-electrode latency as a function of time delay between the index electrode and all other constituent electrodes in each propagation signal. We used the well-known temperature sensitivity of action potential propagation (Chapman, 1967; Franz and Iggo, 1968; Westerfield et al., 1978) to tested the ability of this method to resolve small changes in propagation velocity. The map of MEA electrodes with super- and sub-threshold eAPs of the propagation signal in this example is shown (Figure 3C, top). Recordings were done at the three indicated temperatures and the eAP latency between the index electrode (B7) and other constituent electrodes is seen in averaged spikes from a subset of these electrodes (Figure 3C, below). Superimposed black, grey and red sweeps are averaged, peak-scaled eAPs at the indicated temperatures and show that, as expected, increasing the temperature decreases the eAP latency between the index electrode and each of the other electrodes. We measured the temperature-induced changes in action potential propagation by plotting the change in eAP latency as a function of the propagation time in control conditions (30° C) for every constituent electrode of this propagation signal (Figure 3D, E). The slope of the fitted line from the comparison of two back-to-back recordings done at 30° C was essentially flat (m = 0.004 ± 0.004; Figure 3E). As expected, however, increasing the temperature decreased the latencies between eAPs at the index electrode (B7) and all other electrodes and increased the slopes of the fitted data. For example, raising the temperature from 30° C to 33° C increased the slope of the fitted data to 0.147 ± 0.005, resulting in a 16.2% increase in propagation velocity. Increasing the temperature to 36° C further decreased the inter-electrode latency and increased the slope of the fitted data (0.246 ± 0.007) reflecting that the action potential propagation velocity was 29.9% faster at 36° C compared to 30° C. The fits of these data reflect the linearity of action potential propagation at this level of spatial and temporal resolution. These temperature-induced changes in latency from 15 propagation signals from 3 different MEAs show the robustness of this method (Figure 3F). These data show that this method of quantifying the effects of manipulating propagation velocity easily resolves the effect of small latency differences at multiple sites in axonal arbor and circumvents the need to know the propagation path length.

### Axons in cultured neurons are expansive and contain multiple branches

The two-dimensional extent of constituent propagation signal electrodes is large but is well within the range expected from the reported length and branching patterns of hippocampal axons (Arszovszki et al., 2014; Kaech and Banker, 2006). We validated the extent of axonal growth in our culture system by immunostaining cultured neurons with a marker of neuronal processes (Tuj1) and the presynaptic marker VGluT1, a marker of presynaptic terminals. The plating density of neurons for this experiment was 20% of the MEA plating density to increase our chances of visualizing single axons with minimal contamination from processes belonging to other neurons. Axons were morphologically differentiated from dendrites by their caliber, longer length and more gradual taper (Kaech and Banker, 2006). Figure 4 shows a single neuron with an axon that bifurcates part way along its length. The axonal branches span hundreds of microns and have meandering paths. The areas in the left center and lower right are where branches from this axon enter regions with relatively high levels of VGluT1 staining and large increases in the complexity of processes from other neurons. This axon also showed several small branches off the main bifurcations; the inset highlights a portion of the axon where two smaller axonal branches depart from the main fork of the branch on the left. This example shows that axons in this culture system have meandering paths and extensive branching, consistent with the large and varied patterns of propagation signal electrodes in our system. This validates that axonal processes could represent the physical basis of propagation signals in our recordings.

**Figure 4.**
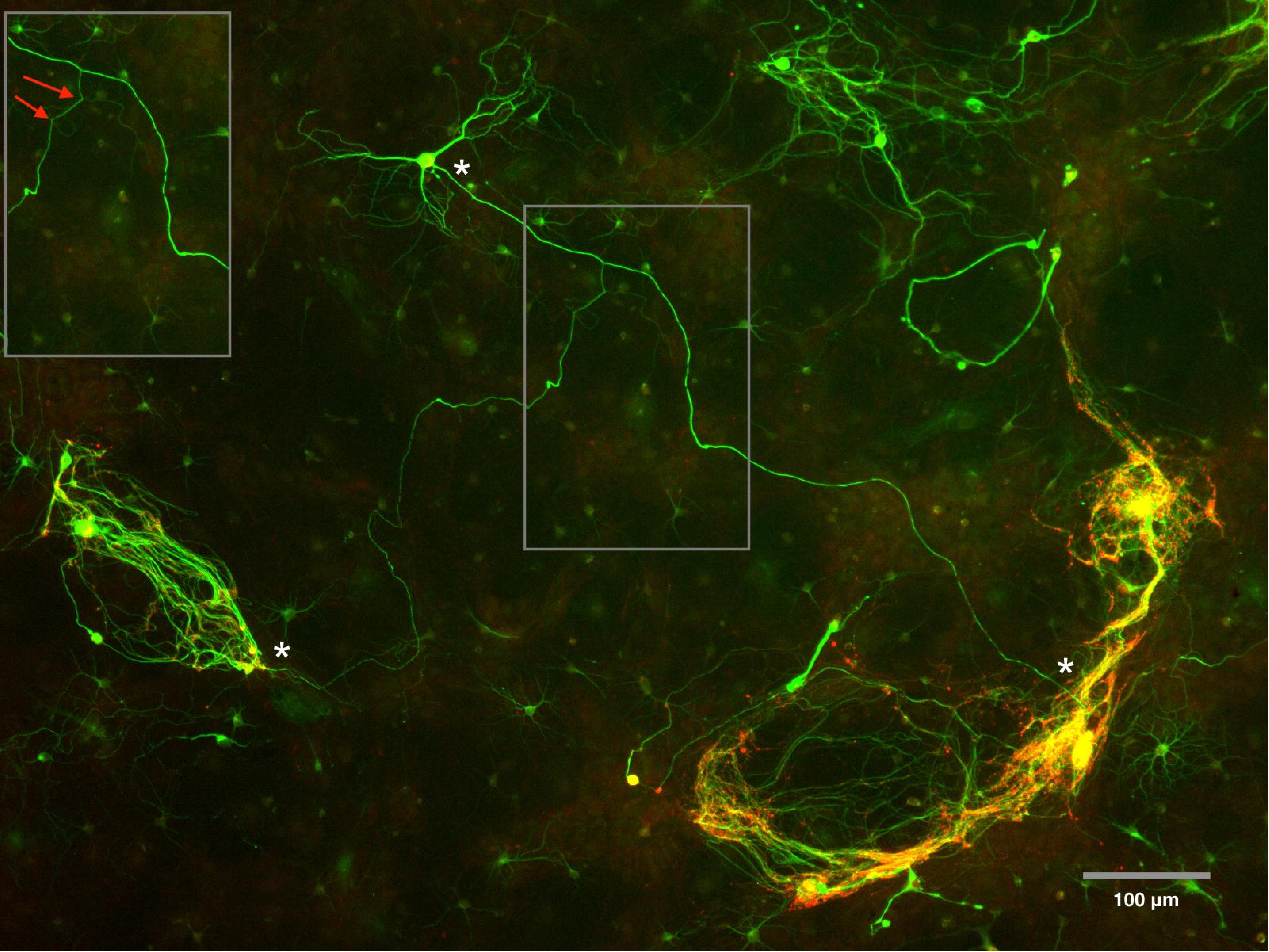
The spatial patterns of electrodes that detect propagation signal components can span hundreds of microns and contain several branches. To examine whether axons displayed comparable morphology to the morphology inferred from propagation signals, we immuno-stained hippocampal neurons with a marker expressed in axons and dendrites (Tuj1) in low density cultures. In these cultures, axons are differentiated from dendrites by their uniform caliber, longer length and more gradual taper. Axons from these cultured neurons can span many hundreds of microns, have meandering paths and several branches. The total length of the process shown between the asterisks was 1692 μm. These images confirm that neuronal processes such as axons could underlie the physical basis of propagation signals in our recordings.

## Propagation Signals are from Axons

Axons have many times the density of voltage-gated sodium channels compared to dendritic or somatic membrane (Hu et al., 2009; Lorincz and Nusser, 2010). Our routine detection of action potential propagation suggests that these eAPs originate from neuronal compartments with a high transmembrane conductance like axons. However, we cannot exclude the possibility that some of these signals result from action potential back-propagation into somato-dendritic cellular regions (Stuart and Sakmann, 1994; Spruston et al., 1995). The large voltage-gated sodium channel density gradient between these compartments is expected to affect the safety factor of action potential propagation (Tasaki, 1953) as has been reported (Mackenzie and Murphy, 1998). Therefore, decreasing the density of active sodium channels with sub-saturating concentrations of the voltage-gated sodium channel blocker tetrodotoxin (TTX) would collapse the propagation safety factor in dendrites to a greater extent than the axonal safety factor. Neuronal sites with low safety factor would experience action potential propagation failures and thus we would expect a dose-dependent change in mean eAP amplitude at these neuronal sites.

The raw voltage traces in Figure 5A show in the same electrode that the amplitude of control eAPs (left) were not noticeably reduced by 10 nM TTX (right). The electrode maps indicate the electrodes at which we detected super- or sub-threshold eAPs (black) for two different propagation signals (Figure 5B). As seen from the averaged waveforms, neither the super- nor sub-threshold eAPs amplitudes were attenuated by 10 nM TTX. Measurement of eAPs from several propagation signals demonstrate that TTX failed to consistently reduce the mean amplitude of super- or sub-threshold eAPs propagation signal components (Figure 5C). The Na_V_1.6 (Scn8a) sodium channel subtype is highly expressed in hippocampal principal neurons (Schaller and Caldwell, 2000). Based on the equilibrium dissociation constant of 6 nM for these channels (Smith et al., 1998), 10 nM TTX is expected to reduce the density of active sodium channels by more than 60%. Thus decreasing the density of active sodium channels by 60% had no effect on the amplitudes of propagation signal eAPs, consistent with propagation signal components originating from neuronal membrane with a high safety factor like axons.

**Figure 5.**
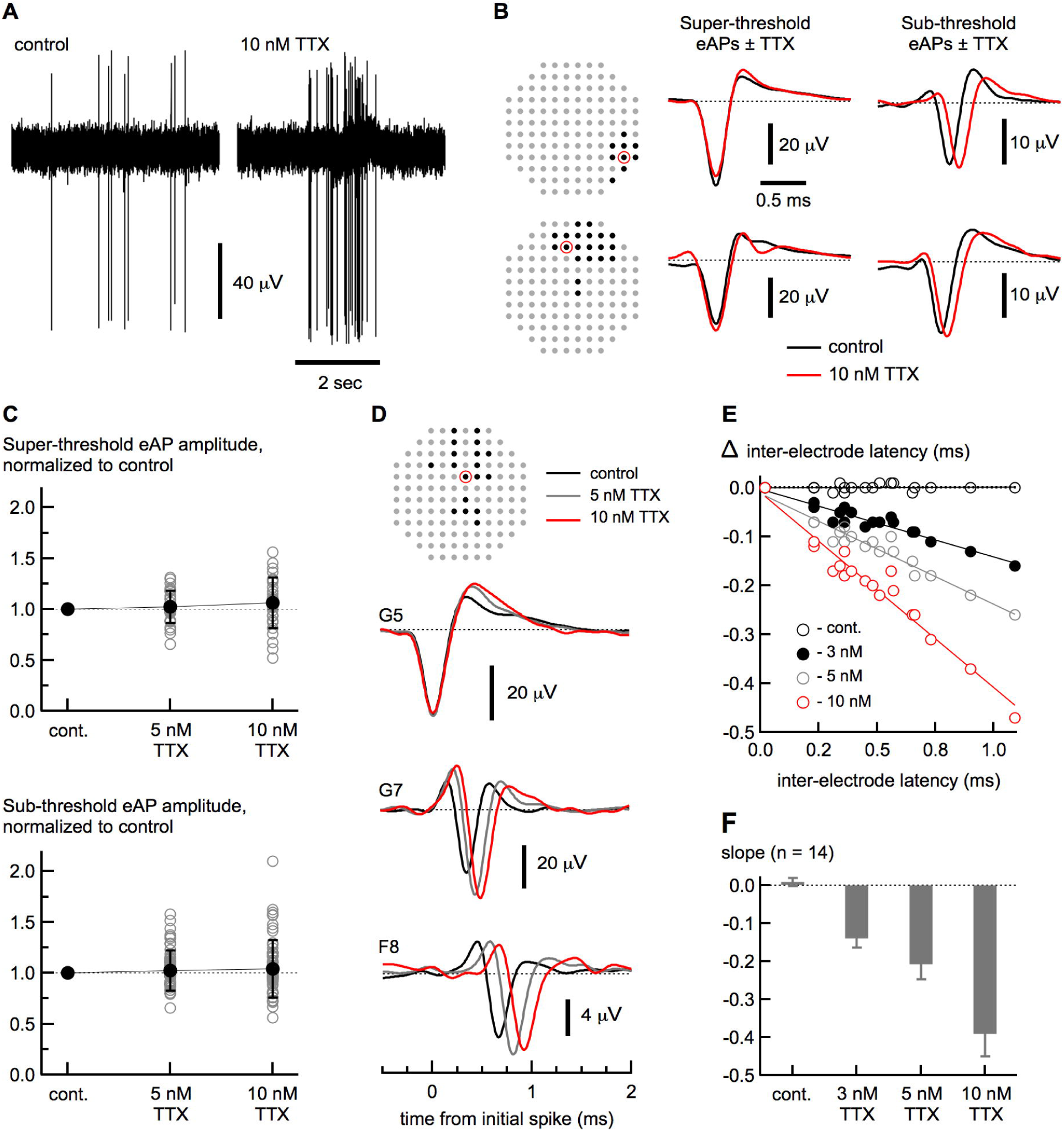
Propagation signals measure axonal action potentials. Extracellular voltage traces from a single electrode are shown (A) in control conditions (left) and in 10 nM TTX (right). Signal-averaged spikes from two different propagation signals (array maps on left) are shown (B), with super- threshold (center) and sub-threshold (right) spikes indicated. Control spikes (black traces) and spikes in TTX (red traces) are superimposed. The mean amplitudes of super- and sub-threshold spikes (normalized to control) from all the propagations signals in these recordings is shown (C) superimposed on individual data points for these conditions. 10 nM TTX had no effect on spike amplitude in either super-threshold spike amplitude (1.06 ± 0.25 compared to control; n = 29 electrodes) or sub-threshold spike amplitude (1.04 ± 0.28 compared to control; n = 67 electrodes). These data are consistent with propagation signals predominantly representing action potential in axons. Data are from from 8 propagation signals on 2 MEAs. Dashed lines are the normalized amplitude of control spikes. In another propagation signal from these experiments (D), TTX decreased propagation speed, as expected from decreasing the density of active sodium channels. Figure 4D shows spikes from a subset of electrodes from this propagation signal, with control spikes (black), spikes in 5 nM TTX (grey) and 10 nM TTX (red) superimposed. Traces in 3 nM TTX were excluded for clarity. In each panel, the dose-dependent increase in latency is seen with the rightward shift of the spikes in increasing TTX concentration. A plot of the change in latency in each condition (control, 3, 5 and 10 nM TTX) as a function of the latency between the initial spike and each spike time in control conditions is shown (E). Data for each condition were fitted with a straight line function. The increasingly negative slope with decreasing active sodium channel density is consistent with a dose-dependent slowing of propagation velocity (F). Data are presented as mean ± SD. Red circles in B and D indicate the index electrode for the indicated propagation signal.

A high safety factor ensures that action potentials occurring at high frequency are faithfully propagated. This is because at higher frequency, voltage-gated sodium channels accumulate in inactivated states (Goldin, 2003). As seen in the eAPs (Figure 5A) sub-saturating concentrations of TTX coincidentally increased the eAP firing frequency and in these cultures. TTX application (3 - 10 nM) changed the spiking pattern and produced array-wide bursting. The median inter-spike intervals (ISI) across all MEA electrodes in the absence of TTX were 211.1 ms and 78.6 ms. These numbers decreased to 21.9 ms and 19.0 ms in 10 nM TTX, respectively. The burstiness indices (Kumbhare and Baron, 2015), a measure of spike distribution across all electrodes, were lower in the control samples (0.456 and 0.602) than in 10 nM TTX (0.929 and 0.976), reflecting array-wide bursting. For propagation signals from individual neurons in control conditions, the average median ISI was 383.0 ± 611.2 ms (n = 19). In the presence of TTX, the average median ISI decreased to 27.5 ± 11.5 ms (5 nM; n = 19) and 35.6 ± 23.1 ms (10 nM; n = 19 neurons). The number of spikes in 20 nM TTX (>95% of control) was insufficient for comparative statistical analysis. The increase in firing frequency caused by low concentrations of TTX made this a rigorous test of how reducing the safety factor affects the fidelity of action potential propagation across the two-dimensional extent of our sampling area. Even at increased firing frequencies (∼30 Hz), reducing the propagation safety factor by TTX application did not reduce the mean amplitudes of super- or sub-threshold eAPs.

Experimentally decreasing sodium channel density decreases axonal action potential propagation velocity (Colquhoun and Ritchie, 1972; Hu and Jonas, 2014). In our experiments, TTX increased the latency between constituent propagation signal electrodes in a dose-dependent manner (Figure 5D). Note the increase in latency between the index electrode (G5) and electrode F8 in 10 nM TTX, for example (Figure 5D; red traces). Even lower TTX concentrations (3 and 5 nM) significantly increased the inter-electrode latency compared to control (Figure 5E-F). Dose-dependent increases in propagation time in these low TTX concentrations are easily resolved by fitting the change in propagation time in TTX as a function of the control propagation time for each propagation signal (Figure 5E). The average slope of the fit for each condition were distinct for each TTX concentration tested (Figure 5F); the mean slopes from fits to the data were 0.01 ± 0.01 in the absence of TTX, -0.14 ± 0.02 in 3 nM, -0.21 ± 0.04 in 5 nM and -0.39 ± 0.06 in 10 nM TTX. These fits reflect decreases in action potential propagation velocity of 12.2%, 17.1 % and 28.1% caused by 3, 5 and 10 nM TTX respectively. Thus under the firing frequencies we report here (∼30 Hz) even when the density of active voltage-gated sodium channels is predicted to be reduced by more than 60% at the highest TTX concentration we tested, axonal action potential propagation still occurs with high fidelity. The high safety factor may be a general property of these types of unmyelinated axons and may mitigate other features affecting propagation fidelity such as inhomogeneity in electrical capacity resulting from extensive *en passant* synapses and axonal branches (Lüscher and Shiner, 1990a; Lüscher and Shiner, 1990b; Mainen et al., 1995). That these TTX concentrations increased inter-electrode latency without affecting the fidelity of action potential propagation indicates an unexpectedly high density of voltage-dependent sodium channels.

## Time-dependent changes in axonal excitability

The non-invasive nature of extracellular recording is well suited to long-term monitoring of neuronal activity and propagation signal eAPs unequivocally identify them as originating from single neurons. Thus propagation signals allow us to monitor excitability in the axonal arbors of individual neurons over time scales from hours to days. For example, the constituent electrodes of one example propagation signal we monitored during a 25-hour period are shown (Figure 5A), with traces from a subset of electrodes from this signal (Figure 5B). The amplitudes of the majority of eAPs at constituent electrodes from this signal were quite stable (eAPs at K5 and F3, for example) during the recording interval. In the same axonal arbor, however, we recorded the *de novo* emergence of an eAP (H4) and the disappearance of another (E2) over the same time period. The waveforms at electrode H4 show the downward (resistive) component of the spike develops from the pre-existing upward (capacitive) component, consistent with the gradual and progressive appearance of active conductances associated with the transmembrane action potential within an existing portion of axon. Thus within 14 hours, the initially non-excitable segment of axonal membrane became invested with enough voltage-dependent conductances to generate a detectable signal in this example. In the case of eAP elimination at electrode E2, the time-dependent decrease in capacitive and resistive eAP components were comparable, consistent with physical elimination of the axonal process rather than a decrease in spike-coupled trans-membrane current in the axonal membrane at that electrode.

The properties of propagation signals allow us to unambiguously identify and monitor single neurons over multiple days. The electrode map (Figure 5C) shows constituent electrodes from one propagation signal that we monitored for 15 days. The earliest occurring spike of this signal was D6, with propagation extending outward diagonally in both directions. The electrode maps of this propagation signal illustrate expansion and contraction of excitability of different portions of the axonal arbor of a single neuron over time. The number of constituent electrodes that sample eAPs from this axonal arbor are shown over time (Figure 5E). Within the same monitoring window, we detected 68 unique propagation signals from 4 MEAs. The distribution of the total number of days each of these signals was detected seemed bimodal (Figure 5F), with many (18) signals being detected throughout the 15 day time window, and almost as many (17) being detected only on a single day. We never recorded a case in which a propagation signal disappeared, only to re-appear at a later time.

Because propagation signals fingerprint the source neuron, we can non-invasively record time-dependent changes in individual eAP waveforms within each propagation signal over long time periods. For example, the eAP at D6 always preceded eAPs at all other electrodes of this propagation signal throughout the 15 day recording period (Figure 5C). The time-dependent increase in spike amplitude at D6 (Figure 5D), coupled with an eAP waveform lacking an initial upward capacitive peak (Figure 5G) is consistent with large increases in transmembrane current at or near the AIS. Additionally, the peak-to-peak interval of the spike waveform at D6 gradually shortens (Figure 5G, top). The peak-to-peak interval of the extracellular eAP approximates the width at half height of the transmembrane action potential (Bean, 2007). Thus the eAP waveform reflects a time-dependent decreases in the transmembrane action potential duration at D6. At the majority of electrodes over 15 days of non-invasive monitoring, the latency between spikes at D6 and eAPs at a subset of electrodes (C8, D5 and G8) initially showed a decrease in the inter-electrode latency between day 11 and day 17, followed by an increase thereafter. In contrast, the time delay between spikes at D6 and F11 also showed a time-dependent decrease but remained stable after day 17 (Figure 5H). These observations indicate that different portions of the axonal arbor from single neurons can act independently from other portions and illustrate the heterogeneity of time-dependent changes in propagation speed across axonal arbor components in single neurons.

## Propagation signals in human iPS-derived neurons

Human iPS-derived neurons are an attractive system in which to study the effects of naturally-occurring mutations, such as those found in sodium channel genes from patients with several forms of epilepsy (Catterall, 2014). Various aspects of action potential propagation in cells from these patients could be assessed with the methods we outlined. For these reasons, we examined whether we could detect propagation signals in human iPS-derived neurons. We reliably detected propagation signals in most cultures of iPS-derived cultures we examined. Figure 6A shows array maps and two propagation signals from the same MEA. A subset of component propagation signal eAPs shown below (Figure 6B). Under the differentiation and cell culture conditions used for these cells, spikes tended to be smaller than spikes recorded in cultured hippocampal neurons but almost all the MEA records we examined had at least one obvious propagation signal. This indicates that human iPSC-derived neurons can be used to study human diseases that affect action potential propagation and the development of axonal excitability. Extracellular recording of propagation signals could make the study of axonal physiology routine.

**Figure 6.**
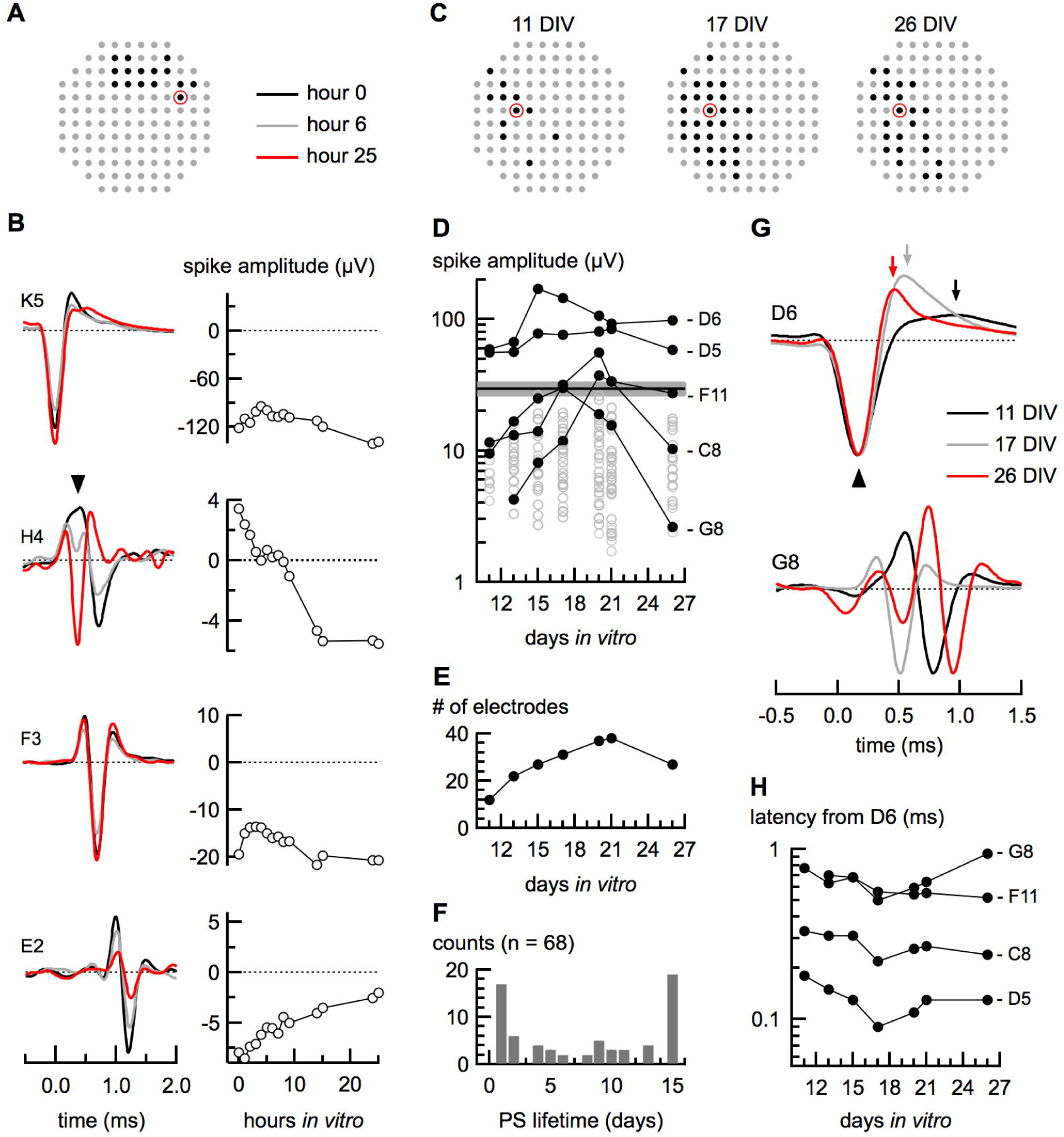
Non-invasive monitoring of axonal excitability during neuronal network development. (A) shows the electrode map of one propagation signal that was monitored over the course of 25 hours with eAPs from a subset of these electrodes shown below (B). Stable eAPs (at K5, F3) were measured simultaneous with the appearance of spikes (H4) and the elimination of spikes (E2) indicating that these processes can occur within the same axonal arbor. Signal amplitudes for each waveform are indicated at right. A different propagation signal was monitored for 15 days (C). The electrode maps show the constituent electrodes at each recording time. Spike amplitudes from all contributing electrodes are plotted as a function of days *in vitro*, with a subset of these is highlighted (black line), indicating time-dependent changes in signal amplitude (D). The grey band is the mean threshold for all electrodes at 21 days in vitro. The total number of constituent electrodes for this propagation signal at every time point is plotted (E). The distribution of propagation signal lifetimes is plotted for 68 propagation signals from four MEAs that we monitored across 15 days (F). The spikes at electrode D6 are the earliest occurring for this propagation signal sequence (from example in C, D) and have a waveform consistent with being at or near the site of action potential initiation (G, above). We noted a developmental decrease in spike width of spike at D6, shown by the decrease in the peak-to-peak time from 11 to 20 DIV (arrows). We also noted that a time-dependent change in the timing delays between eAPs at D6 and other electrodes (G, H). The timing difference between the eAP at D6 and the eAP at G8 initially decreased then increased to a latency greater than seen in the first measurement (G, below). Changes in spike latencies were non-uniform across the axonal arbor (H). Red circles in A and C indicate the index electrode for the indicated propagation signal.

**Figure 7.**
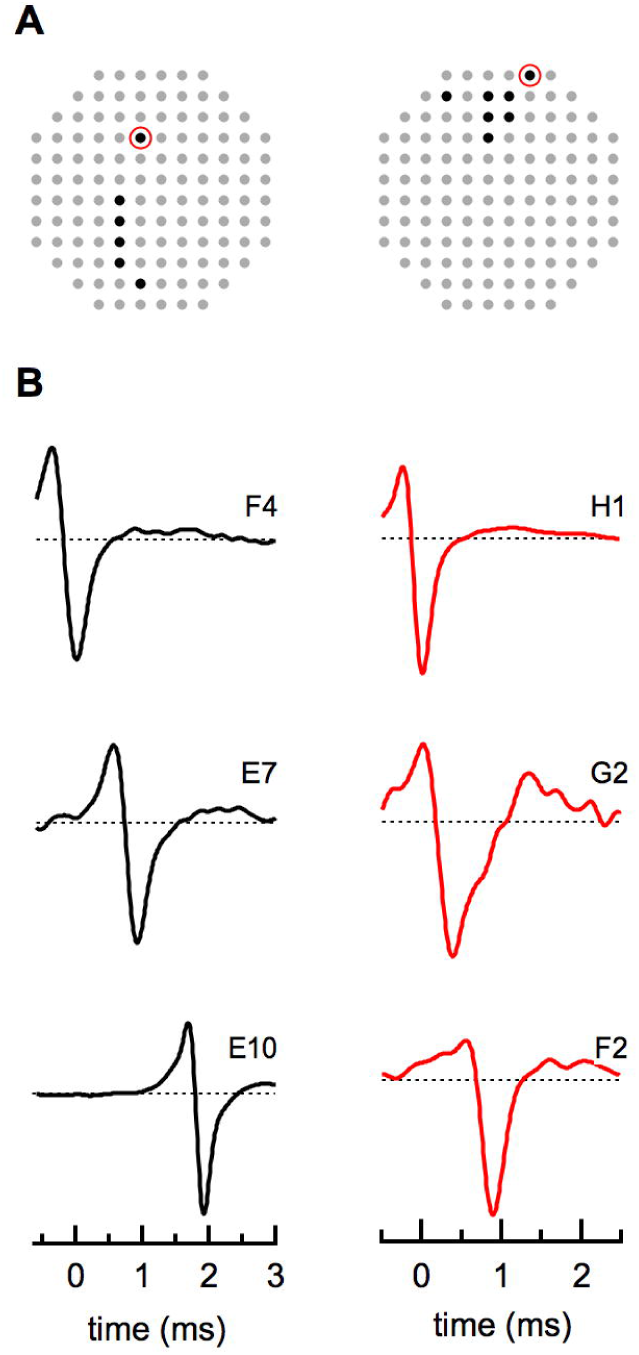
Propagation signals from human IPSC-derived neurons. The electrode map (A) shows the electrode positions of two propagation signals from the same array. Example eAPs from a subset of these electrodes is shown (B). Amplitudes of the eAPs were peak-scaled to the largest spike amplitude within each propagation signal.

## Discussion

Axons are responsible for the integration and high fidelity transmission of neuronal input (Debanne et al., 2011). However, due to their small caliber, axons are a privileged neuronal compartment from the perspective of routine experimental accessibility. The data presented here show that low electrode density MEAs can be used to routinely assess action potential propagation from single neurons within neuronal networks. Repeated co-occurrence of eAPs on multiple electrodes and the invariability of the timing sequence between those electrodes is a fingerprint that unambiguously validates that these signals result from single neurons. In the context of the extracellular recording configuration, it is unprecedented to routinely and unambiguously determine that a subset of eAPs results from a single neuronal source in a background of eAPs from other cells. Because these eAPs come from a known neuronal source, averaging these signals reveals the greater extent over which we can record eAPs in the axonal arbor and increases the range of time differences used for assessment of the effects of manipulating propagation. Signal averaging also allows us to resolve biophysical-level details of spikes from different parts of the axonal membrane. The ability to ‘fingerprint’ enables us to follow individual neurons within the greater network over several days. Finally, propagation signals in human iPSC-derived neurons could be used to study axonal physiology within the context of human disease models.

MEAs have previously been used to measure action potential propagation in axons. However, this work was done using custom designed arrays with thousands of electrodes (Bakkum et al., 2013; Müller et al., 2015) or in micro-fluidic chambers designed to constrain axonal growth (Dwowak and Wheeler, 2009) and which prevent interaction with other neurons. The large number of electrodes used in the former technique can produce very large data files that may limit the high-throughput use of this technology as well as limit the increased signal to noise obtained by averaging many hundreds of waveforms obtainable in a longer recording. The latter technique necessarily isolates axons from greater network interaction. Because axons of many hippocampal neurons have *en passant* presynaptic terminals along their length, isolating axons in this way could mask the contribution of presynaptic conductances during action potentials and membrane inhomogeneity on action potential propagation, for example. Large-scale MEAs have also been used to measure propagation in the retina (Li et al., 2015) in experiments that coupled electrical measurements with fluorescent labelling of small numbers of retinal ganglion cells. This combined approach of fluorescent labelling with electrical measurement is attractive but impractical to implement in high density neuronal cultures because fluorescently labelling all neurons makes following single processes impractical. Labelling a fraction of neurons may routinely fail to label neurons that underlie propagation signals. One technique couples fluorescent labelling with electrophysiological recording and thus could be used to label single identified propagation signal neurons in our system (Pinault, 1996). However, our analyses rely on changes in propagation time between component electrodes of each propagation signal (Figure 3) and thus circumvent the need to know axonal path length. Thus while our methods do not give absolute values for propagation velocity, our ability to non-invasively sample from multiple sites provides the dynamic range to quantitatively study the effects of manipulating propagation. This creates avenues for the routine study of the physiological and pathophysiological parameters affecting axonal action potential propagation.

## Signal redundancy

Propagations signals were found in every recording done in mouse neuronal cultures and our data directly demonstrate that eliminating propagation signals on all but one constituent electrode reduced the total number of spikes by 40%. Propagation signals affected, on average, 25% of the electrodes in arrays such as the ones we used in these experiments. Doubling the electrode spacing can minimize the contribution of simultaneous spike detection between neighboring electrodes but does not eliminate the redundancy resulting from propagation signals. It is surprising that this form of signal redundancy has not been more commonly reported. This could be due to the use of larger pitch electrodes by other investigators (Shahaf and Marom, 2001; Wagenaar et al., 2005), potentially decreasing the likelihood of detecting propagation signals.

The prevalence of propagation signals in our data records means that valid assessment of the significance of array-wide behavior requires their elimination. Spike sorting routines attempt to assign spikes recorded on individual electrodes into units, for the purpose of further spike train analysis (Hill et al, 2011). Detection and elimination of propagation signals from all but one recording channel is itself a novel spike sorting step and may in fact make subsequent algorithm-based sorting more robust. Because propagation signals result from the same neuron they are a source of ground truth that can be used for testing spike sorting algorithms (Quiroga, 2013). Our catalog of propagation signals is a resource to examine signal heterogeneity from single neurons.

Our cell culture technique could also account for why we readily observe propagation signals. Glia from dissociated hippocampi are grown to confluence and only then does a second round of plating occur meaning that neurons grow on a preexisting glial substrate. This could create a permissive environment for growth of axons closer to MEA electrodes than they might otherwise grow. Alternatively, glia could act as a resistive sheet that increases eAP amplitude across all electrodes (Matsumura et al., 2016). On bare arrays we observed that neuronal processes tend to avoid the area around the electrodes, that the signal-to-noise ratio was lower than in neurons plated on a glial bed and propagation signals were not as notable (data not shown). It is also possible that glial/neuron apposition increases the transmembrane current density in neurons through glial-neuronal signaling mechanism (Tang et al., 2014; Sobieski et al., 2015; Fields 2015) or by increasing the axon length and complexity of branching (Hughes et al., 2010).

## Future Directions

Axons and the axon initial segment are considered important final mediators in conveying signals to postsynaptic partners (Kole and Stuart, 2012; Debanne et al., 2011). However, techniques used to study the physiological properties of axons, such as patch clamp-based recording configurations are technically challenging (Schmidt-Hieber et al., 2008; Casale et al., 2015). We have shown that comparable or parallel information can be routinely obtained from extracellular recordings with MEAs. Moreover, unlike recording in the patch clamp configuration, extracellular recording is non-invasive, meaning that the spiking behavior from single neurons can be monitored across hours and days. As we demonstrated, the development of excitability in axons proceeds gradually; expansion and retraction can occur simultaneously in different parts of the same axonal arbor (Figure 6B) and thus elaboration of the axonal arbor is not necessarily a concerted process. The earliest propagation signal eAP is the index around which our measurements occur because thess spikes tends to be the largest and often have properties consistent with eAPs that originate from at or near the AIS. These features create a basis from which to explore the development of axonal excitability.

The high-throughput nature of extracellular recording with MEAs means that propagation signals can also be used to study channelopathies of ion channels that are localized to axons and to ask how these or other mutations affect propagation velocity. Mutations have been found in several types of voltage-gated ion channels that are typically localized to axons and are associated with phenotypes such as congenital epilepsy as well as neurodevelopmental disorders such as autism (Debanne et al., 2011; Kullman, 2010). The effects of these mutations on axonal excitability could be assessed with the approaches we have outlined. Our methods could, for example, be used to study the effects of temperature on action potential propagation in models of Dravet syndrome, a condition in which patients have a high incidence of febrile seizures and which, in the majority of cases, is associated with mutations in the Scn1a subtype sodium channel (Catterall, 2014). Propagation signals could be used to study other mutations in Scn1a that are associated with other seizure phenotypes. Patients with polymorphisms at Scn1a can also vary in their responsiveness to commonly prescribed anti-epileptic treatments (Tate et al., 2005). Propagation signals could be used to understand the biophysical nature of these differences in drug sensitivity in patients with these polymorphisms.

## Acknowledgements

This research was sponsored by the U.S. Army Research Laboratory and Defense Advanced Research Projects Agency under Cooperative Agreement Number W911NF-15-2-0056. The views, opinions, and/or findings contained in this material are those of the authors and should not be interpreted as representing the official views or policies of the Department of Defense or the U.S. Government. Additional support was also provided by the California NanoSystems Institute (CNSI). We thank Bridget N. Queenan and Carol A. Vandenburg for their thoughtful reading and insightful comments on this manuscript.

